# Fine-tuned control of stress priming and thermotolerance

**DOI:** 10.1101/2021.01.27.428215

**Authors:** Benjamin Pfeuty, Emmanuel Courtade, Quentin Thommen

**Affiliations:** Univ. Lille, CNRS, UMR 8523 - PhLAM - Physique des Lasers Atomes et Molécules, F-59000 Lille, France

## Abstract

A common signature of cell adaptation to stress is the improved resistance upon priming by prior stress exposure. In the context of hyperthermia, priming or preconditioning with sublethal heat shock can be a useful tool to confer thermotolerance and competitive advantage to cells. In the present study, we develop a data-driven modeling framework that is simple and generic enough to capture a broad set of adaptation behaviors to heat stress at both molecular and cellular levels. The model recovers the main features of thermotolerance and clarifies the tradeoff principles which maximize the thermotolerance effect. It therefore provides an effective predictive tool to design preconditioning and fractionation hyperthermia protocols for therapeutic purpose.

## 1. Introduction

Living cells are exposed to environmental stressors that fluctuate and repeat over a wide range of amplitude and time scales. The efficiency and speed of cellular response and adaptation are highly dependent on the well-regulated dynamics of their signaling and regulatory network [1, 2, 3]. The diverse stress response pathways involve specific proteins and mechanisms but share nevertheless a similar architecture where environmental stress induces intracellular damages but also upregulates damage repair pathways and programed cell death pathways [4, 5].

The search for optimized stress protocols taking advantage of Achilles’ heels of cells [6] is a quest for decades especially in therapeutic field. The initial approach based on a dual control of intensity and duration of the stressor while measuring the surviving cell fraction to establish dosimetry tools [7, 8, 9], has been paralleled by sensitization and fractionation strategies [10, 11]. The effects of fractionation are in fact widely used in the treatment of cancer by radiotherapy. Cellular adaptation to stress has been described in the available dosimetry standards based on regression, for lack of a better term. However, the cellular mechanisms of the underlying regulatory networks have since been identified thanks to sustained experimental efforts. It is therefore tempting to revisit the issue of cellular adaptation to recurring stress with the most comprehensive approach of systems biology [12].

The application of heat in the treatment of malignant tumors has a long history dating back from ancient Greeks. A well-known example dates from the end of the 19th century, when spontaneous regression of malignant tumors was reported in patients with high fevers due to bacterial infections [13]. Nowadays, hyperthermia, also known as thermal therapy or thermotherapy, is still a therapeutic option used either alone or in combination with other chemical or radiative treatments [14, 15]. In addition to direct effects, hyperthermia also appears as a powerful modifier of the tumor response to radiation and several chemotherapy agents because hyperthermia increases and targets their cytotoxic effects in the tumor volume. A multitude of randomized studies has shown that hyperthermia in combination with radiation therapy, chemotherapy or both, resulted in significantly improved clinical outcomes in cancer patients (see [16] for instance). Hyperthermia, associated to temperature ranging from 39 to 45°C, can treat a wide range of tumor types with restricted damage to normal tissue. Tumor sites include cervix, soft tissue sarcoma, breast, head and neck, rectum, brain, bladder, lung, esophagus, liver, appendix, prostate and melanoma [17]. Obtaining effective hyperthermal treatment in the clinic requires both high-quality heating equipment and accurate thermal dosimetry. Current techniques for the application of hyperthermia use various techniques: electromagnetic heating, ultrasound, hyperthermal perfusion and conduction heating, adapted according to the location and extent of the area of the human body to be treated [15].

Despite promising clinical advances, the mechanisms involved in heat-stress-induced cell death have long been debated [18] as hyperthermia causes numerous changes in cells and loss of cell homeostasis [19, 20]. A key event appears to be the denaturation and aggregation of proteins [21] which leads to cell cycle arrest, inactivation of protein synthesis and inhibition of DNA repair processes. The correct structure and conformation of proteins are indeed essential for their function in the cell. A slight increase in temperature can cause unfolding, entanglement and aggregation of proteins, leading to an imbalance in proteostasis. This may result in increased degradation of the aggregated/unfolded proteins through the proteasomal and lysosomal pathways. Other cellular effects of hyperthermia include the inhibition of DNA synthesis, mRNA transcription and protein translation, the disruption of the membrane cytoskeleton, metabolic changes (e.g., decoupling of oxidative phosphorylation) that leads to decreased ATP levels and the alterations in membrane permeability that lead to increased intracellular levels of Na+, H+ and Ca2+ [19, 20].

Thermotolerance is defined as the transient resistance to heat after prior heat treatment occurring in both normal tissue and tumors [22]. Unfortunately, the variation in the kinetics and magnitude of thermotolerance across tissues requires to address cell type-specific characteristics of thermotolerance. The magnitude and kinetics of thermotolerance appear to depend on the thermal damage induced by the priming heat treatment [23]. It is well known that the depletion of the *Heat Shock Protein 70 kDa* (HSP70) molecular chaperone, which is in charge of protein folding and is actively synthesized upon heat exposure, abrogates the protective effects of thermotolerance and sensitizes cells to apoptosis [24, 25].

Recently, a mathematical model of cell survival under hyperthermia has been proposed. In contrast to heuristic thermal dose models, such as the CEM43 [26], this new tool relies on mechanistic modeling of heat stress regulatory network, conferring the possibility to accurately predict cell survival even when the temporal profile of the protocol is complex [27]. The generality of this model has been established by correlating cell line-specific thermal sensitivities with the abundance of molecular chaperones [28]. However, the model still lacks a description of the heat-induced upregulation of molecular chaperone responsible for thermotolerance. Detailed models of the heat-shock transcriptional pathways [29, 30, 31] can be reduced to simple two-variable negative-feedback models when focusing on the transcriptional response at the hour timescale relevant for thermotolerance [32].

Based on these previous modeling investigations, the present study proposes a systematic analysis of thermotolerance at both molecular and cellular levels. Specifically, we address the relationship between the regulated synthesis of molecular chaperones and the thermoprotective effect of a preconditioning treatment. We first develop a model of cell survival to proteotoxic stress including the activation of molecular chaperone synthesis. The model is simulated to study the thermoprotective effect using the standardized measure of ThermoTolerance Ratio (TTR), which quantifies the change in the slope of the survival curves induced by preconditioning treatment. We obtain a resonance-like pattern of the TTR with respect to temperature, indicating the temperature window at which the preconditioning treatment has an optimal thermoprotection. The characteristics of the thermotolerance response is systematically analyzed as function of regulatory and protocol parameters. Such systematic analysis captures the intracellular mechanisms responsible for thermotolerance, but also allows to predict the optimal preconditioning protocols in various setting.

## 2. Results

To study the thermotolerance effect, we combine two dynamical models that have been previously developed to describe, respectively, the transcriptional response [32] and the survival response [28] to a proteotoxic stress, and whose parameters have been estimated from experimental data. The resulting model considers effective relations between proteotoxic damages, molecular chaperone mRNA and protein concentrations and survival rate:

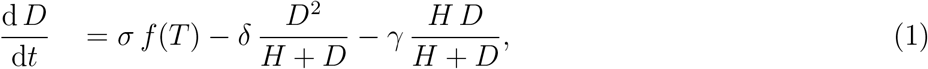

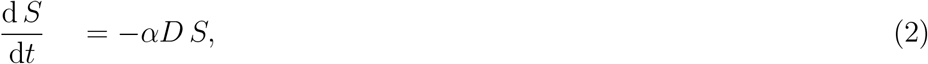

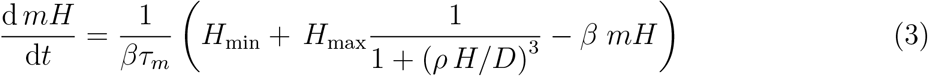

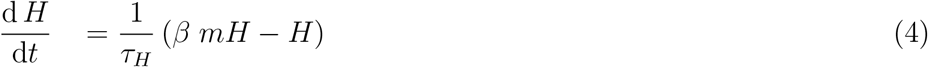

In the first equation, the variable *D* describes the level of damages, here misfolded proteins produced at a temperature-dependent rate *σ f* (*T*) and degraded or renaturated depending on the relative level of damage and molecular chaperones *H*. The function *f* has been previously estimated to 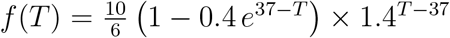 [34]. As well, the relative degradation or renaturation rates of damaged proteins have been demonstrated to depend on chaperone in a simple manner where degradation dominate for *D* ≫ *H* while renaturation prevails for *D* ≪ *H* [27].

The second equation describing survival rate at population level does not include cell division process because the initial value of *S* is set to *S*(0) = exp(*αD*_37_ *t_exp_*) where *D*_37_ is the steady state value of *D* at 37°C and *t*_exp_ defines the experiment duration. Therefore,

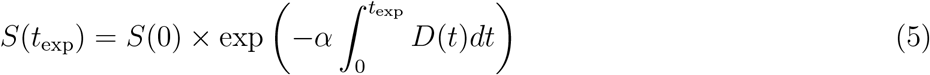

corresponds to the cell survival measured by cell colony assay, where *S*(*t*_exp_) = 1 for a constant exposure at 37°C [28]. In the calculation we set *t*_exp_ = 360 h to mimic the 15 days of rest used in colony formation assay.

In the Equations describing chaperone dynamics, the transcription rate of chaperone mRNA is modeled by a function that ranges from *H*_min_ to *H*_min_ + *H*_max_ depending on the relative abundances between molecular chaperones and damage. Such a dependence of the transcriptional activity corresponds to the chaperone displacement mechanism. While the molecular chaperone HSP70 sequesters HSF1 transcription factor in unstressed basal conditions, the increase of stress and damage leads chaperone to bind damaged proteins thereby releasing HSF1 which further activates the transcription of chaperones. The parameter *ρ* defines the activation threshold of chaperone synthesis relative to the switchover between damage degradation and renaturation (H/D): *ρ* < 1 indicates that the synthesis occurs while renaturation is still the dominant process while ρ > 1 indicates that damage degradation is the dominant pathway.

Most parameters of the model have been estimated from experimental data [28] where *α* = 0.0831 *μ*Mh^−1^, *σ* = 57.4 h^−1^, *δ* = 5.74 h^−1^, *γ* = 1200 h^−1^ have been fixed for all cell lines. RNA and protein half-lives (*τ_m_* = 1 h and *τ_H_* = 30 h) have already been estimated in HeLa cells and are expected to be similar across cell lines [32]. The Hill’s coefficient is set to 3 in relation to the trimerization of HSF1 before transcriptional activation and consistently with experimental measurements [32]. The translation rate is set at a consensus value *β* = 100 *μ*Mh^−1^ keeping in mind that it is a scaling factor for mRNA level that has no dynamical effect. The model contains only three undetermined parameters, *H*_min_ and *H*_max_ which characterize the minimal and maximal molecular chaperone level, and *ρ* which sets the activation threshold. It is convenient to introduce a new parameter denoted *H*_ref_ that represents the level of molecular chaperone *H* at 37°C and which is maintained constant throughout the study. Accordingly, increasing the chaperone regulation strength H_max_ requires to decrease H_min_ in order to keep constant the initial chaperone capacity *H*_ref_. To summarize, the regulatory mechanism involved in the heat stress response is essentially determined by the two free parameters H_max_ and *ρ*, which define respectively the strength and the threshold of the negative-feedback transcriptional control of chaperone synthesis (mediated implicitly by the transcription factor HSF1).

The model described by Eqs 1–4 and parameterized as aforementioned is sufficient to quantitatively capture the main features of cellular adaptation, which are (i) the dynamical overshoot of the molecular response and (ii) the flattening of the survival curve for long enough exposure (Fig. 1). For this purpose, we consider experimental measurement of heat stress response of Jurkat cells, for which the fold change of HSP70 and its mRNA have been recorded upon a 4-hour stress at 41 °C and survival curve is well characterized and shows 10% of cell survival following one-hour heat stress at 43°C [35]. The model adjusts the experimental data for some specific value *H*_max_ = 17.7 *μ*M and *H*_min_ = 0.35 *μ*M. Simulations of the heat shock response for varying values of *H*_max_ (keeping *H*_ref_ = 0.4*μ*M) clearly shows the role of chaperone regulation on both the adaptation profile of the transcriptional dynamics and the flattening of survival curve.

**Figure 1.**
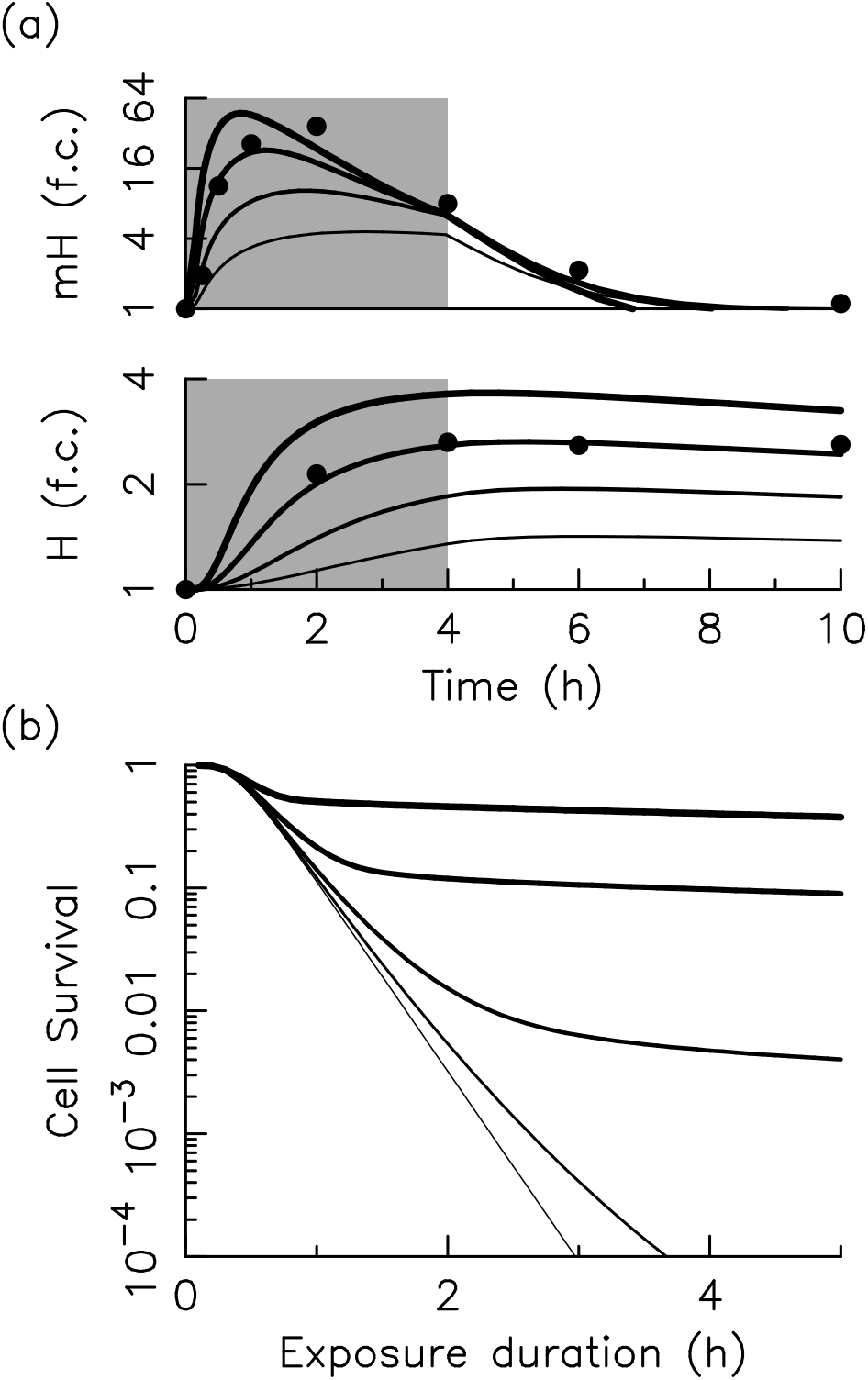
Cellular adaptation to heat stress. (a) Adaptation dynamics of molecular chaperone (mRNA and proteins) induced by a 4-hours heat shock at 41°C in Jurkat cells. Black circles correspond to experimental data from[33]. (b) Survival curves as a function of exposure duration of heat stress. Thicker lines are associated with a stronger regulation (*H*_max_ = 0.4, 1.77, 5.6, 17.7 and 56). The calculation uses a heat stress of 43°C with *H*_ref_ = 0.4μM to show 10% cell survival for a stress duration of one hour (typical value of Jurkat cells).

Fractionation protocols provide additional degrees of freedom and parameters to control the cellular stress response. The most basic fractionation protocol setting is called preconditioning or priming and consists in two stress pulses of respective amplitudes *T*_1,2_ and durations *τ*_1,2_ and separated by a recovery time *τ_r_* (Fig. 2). Cell survival is generally measured with a first stimulus and recovery time of fixed durations, and a second pulse of graded duration, where the effect of priming is evaluated by comparing survival curve with and without recovery. A standard representation depicted in Figure 2 displays the cell survival values in log scales obtained with and without recovery as a function of the total exposure duration (*τ*_1_ + *τ*_2_). A similar representation is used for radiotherapy. The efficiency of the fractionation protocols arising from the recovery phase has long been quantified by the Thermo-Tolerance Ratio (TTR) [22]. TTR is defined as the ratio of the additional durations after priming with and without recovery, respectively named Δ_2_ (*τ_r_ >* 0) and Δ_1_ (*τ_r_* = 0):

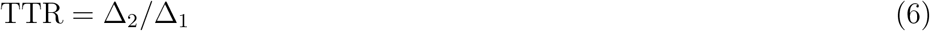

**Figure 2.**
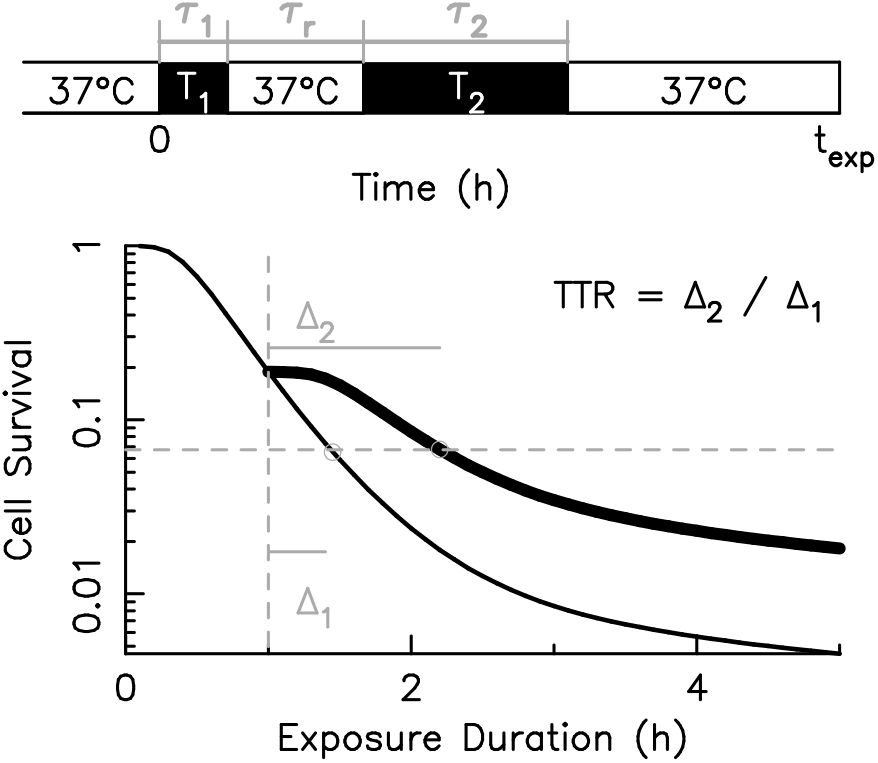
Fractionation protocol and thermotolerance ratio (TTR). A basic fractionation protocol consists in two stress pulses of duration *τ*_1,2_ separated by a recovery period of duration *τ_r_*. The survival curve obtained by varying the *τ*_2_ and represented by the thick line is compared from that obtained without recovery period (*τ_r_* = 0) represented by the thin line. The difference between survival curves is quantified by the TTR defined as the ratio of the additional duration of stress *τ*_2_, TTR = Δ_2_/Δ_1_ (Eq.6).

By defining the following survival function *S* = *f* (*T*, *τ*_1_, *τ*_2_, *τ_r_*), Δ_1/2_ are respectively defined by the implicit relations 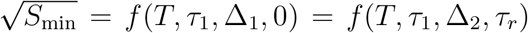. TTR is a dimensionless quantity larger than 1 whose computation only requires to predetermine a reference value of cell survival which is here 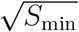 (i.e., half the minimal value at *τ*_2_ = 5h).

The TTR varies strongly with the stress temperature and displays a resonance-like pattern around 43°C (Fig. 3). The thermotolerance phenomenon thus appears in a narrow temperature window. For low proteotoxic stress, the initial chaperone capacity (*H*_ref_) is already sufficient to buffer the low amount of produced damages, and the moderate additional level of chaperone induced by the first stress does not improve significantly the adaptation performance. For a high proteotoxic stress related to temperature above 44°C, the increase of molecular chaperone up to its maximum level after priming remains nevertheless not sufficient to cope with the high level of damages produced during the second heat stress. The thermotolerance therefore arises in an intermediate range of temperature and damage production rate, for which the increase of chaperone induced by the preconditioning stimulus (stress priming) provides an extra-pool of free and functional chaperones that helps to cope efficiently with the damage produced during the second heat stress. The characteristics of such peak of TTR can be described in terms of its amplitude *A_TTR_,* its half-height width between *T*_ and *T*_+_ and the position *T_m_* of its maximum (Fig 3(a)).

**Figure 3.**
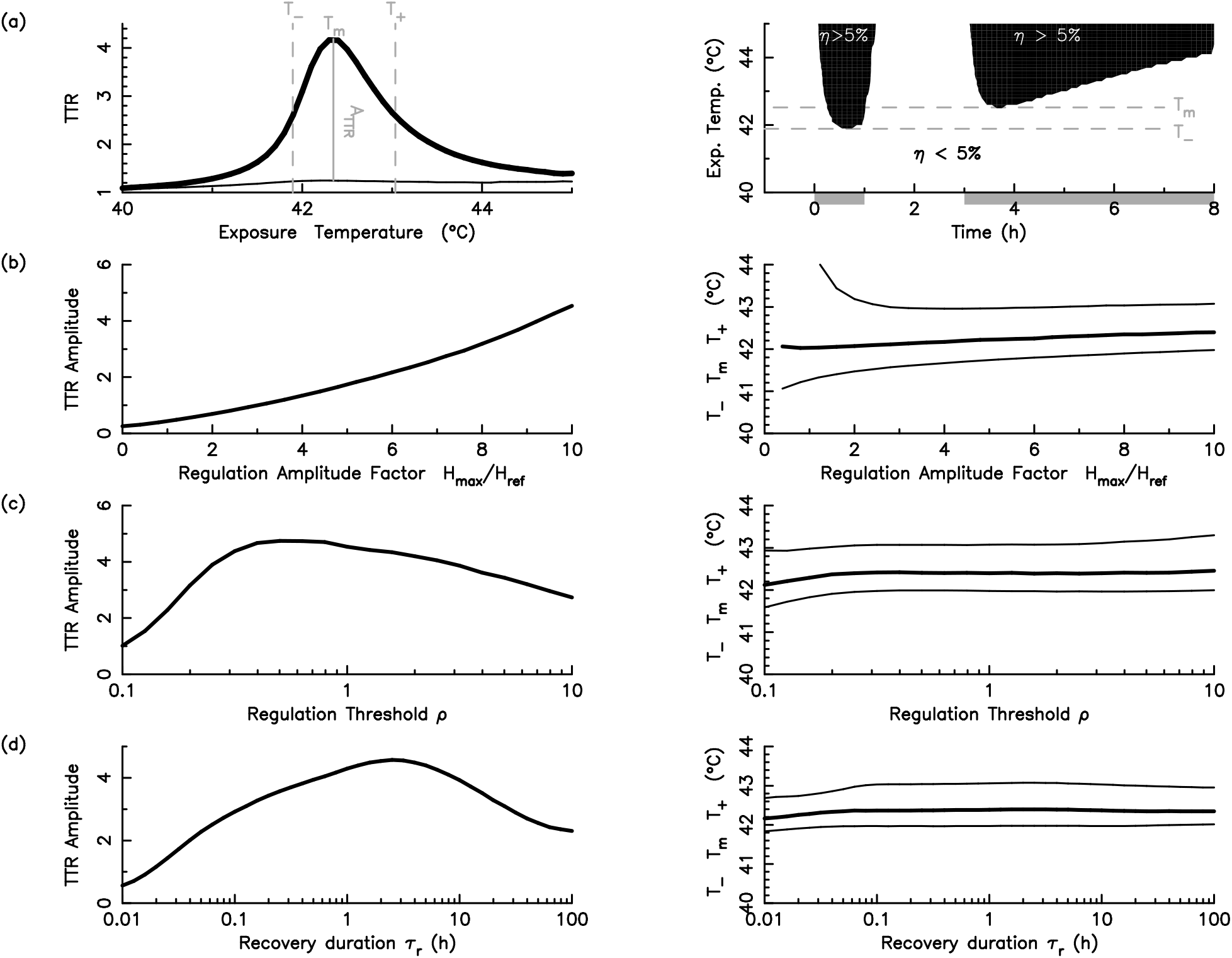
Thermotolerance properties tuned by regulatory and protocol parameters. Parameter values of reference are *τ*_1_ = 1h and *τ_r_* = 2 h for the protocol parameters, *H*_ref_ = 0.4*μ*M, *ρ* and *H*_max_ = 10 *H*_ref_ for the regulatory parameters. (a) The TTR displays a maximum with the exposure temperature around 42.5°C (left panel). The amplitude and position of the maximum, as well as the width of the window, increase with the control force with *H*_max_ = 0 (thin line) or *H*_max_ = 10 *H*_ref_ (thick line). White area on the right part locates renaturation prevalence (*η* < 5%) for various exposure temperature. Gray bar on the bottom indicates thermal protocol (*τ*_1_ = 1 h; *τ_r_* = 2 h; *τ*_2_ = 3 h). (b-d) The maximum TTR value in the range from 40°C to 45°C in the left panels, and the timing of the maximum (thick line) and the half-maximum (thin lines) in the right panels, are computed as function of *H*_max_ (b), *ρ* (c) and *τ_r_* (d), with other parameters taken at their reference values.

Mechanistic understanding of the resonance-like pattern of the TTR relies on the quantification of the fate of misfolded proteins by the triage index *η* = *δ D*/(*δ D* + *γ H*). *η* tends to 0 (resp., 1) when misfolded proteins are preferentially renatured (resp., degraded). As soon as renaturation is close to saturation, typically *η* at 5%, the concentration of misfolded proteins quickly increases and survival sharply decreases. Tracking over time the triage index and for various exposure temperatures of a fractionated protocol indicates that large amplitudes of TTR appear when the triage regimes are different in the two stresses (right panel of Fig 3(a)).

In the following, we perform a systematic analysis of the peak properties of TTR as a function of some key regulatory and protocol parameters (Fig. 3(b)-(d)). As expected, increasing the regulatory strength *H*_max_ gradually amplifies and sharpens the peak while shifting the thermotolerance-effective temperature from 42 to 43°C. In contrast, the TTR amplitude *A_TTR_* displays a maximum for an intermediate value of *ρ*. Indeed, for small values of *ρ*, the transcription function is already at its maximum level in standard condition, and the network has no more adaptive capacity and thus no more possible themotolerance. In turn, for large values of *ρ*, the synthesis is only activated when the degradation process is largely dominant such that increasing the renaturation capacity has no effect. The TTR amplitude also depends significantly on recovery time, with a maximum around 4h and a slow relaxation of timescale similar to the half-life of molecular chaperones. The recovery time has no effect on the position and extent of the peak of TTR.

Besides the recovery time, other properties of the preconditioning protocol can be modulated such as the duration *τ*_1_ or the temperature *T*_1_ of heat stress priming. Mechanistic modeling brings opportunities to predict protocols that would maximize or minimize cell death by enhancing or circumventing the thermotolerance effect. To illustrate the latter case, we consider a lethal hyperthermia treatment of two hours of exposure at 43°C resulting in less than 1% of survival for Jurkat cells (*H*_ref_ = 0.4*μ*M).

Using parameter optimization techniques, we search for the set of protocol parameters that maximize survival as a function of regulatory strength *H*_max_ (Fig. 4). Priming is predicted to raise cell survival from 1% up to 70% for strong regulation (Fig. 4(a)). The optimal heat priming protocol requires rather long initial exposure *τ*_1_ (> 5 h), short recovery time *τ_r_* and a sublethal temperature below 40°C (Fig. 4(b)). It is to note that this optimal priming temperature, which allows to activate the synthesis without penalizing the survival, depends strongly on the regulation threshold *ρ* (*ρ* = 0.3 in Fig. 4). More generally, cell survival significantly varies in the space of protocol parameters (Fig. 4(c)), which can be used to draw optimality manifolds as lines or surfaces. These manifolds can be used to optimally adjust protocols under some particular practical constraints (i.e., the intersection of the optimality manifold and constraint manifold), related for instance with a specific therapeutical context.

**Figure 4.**
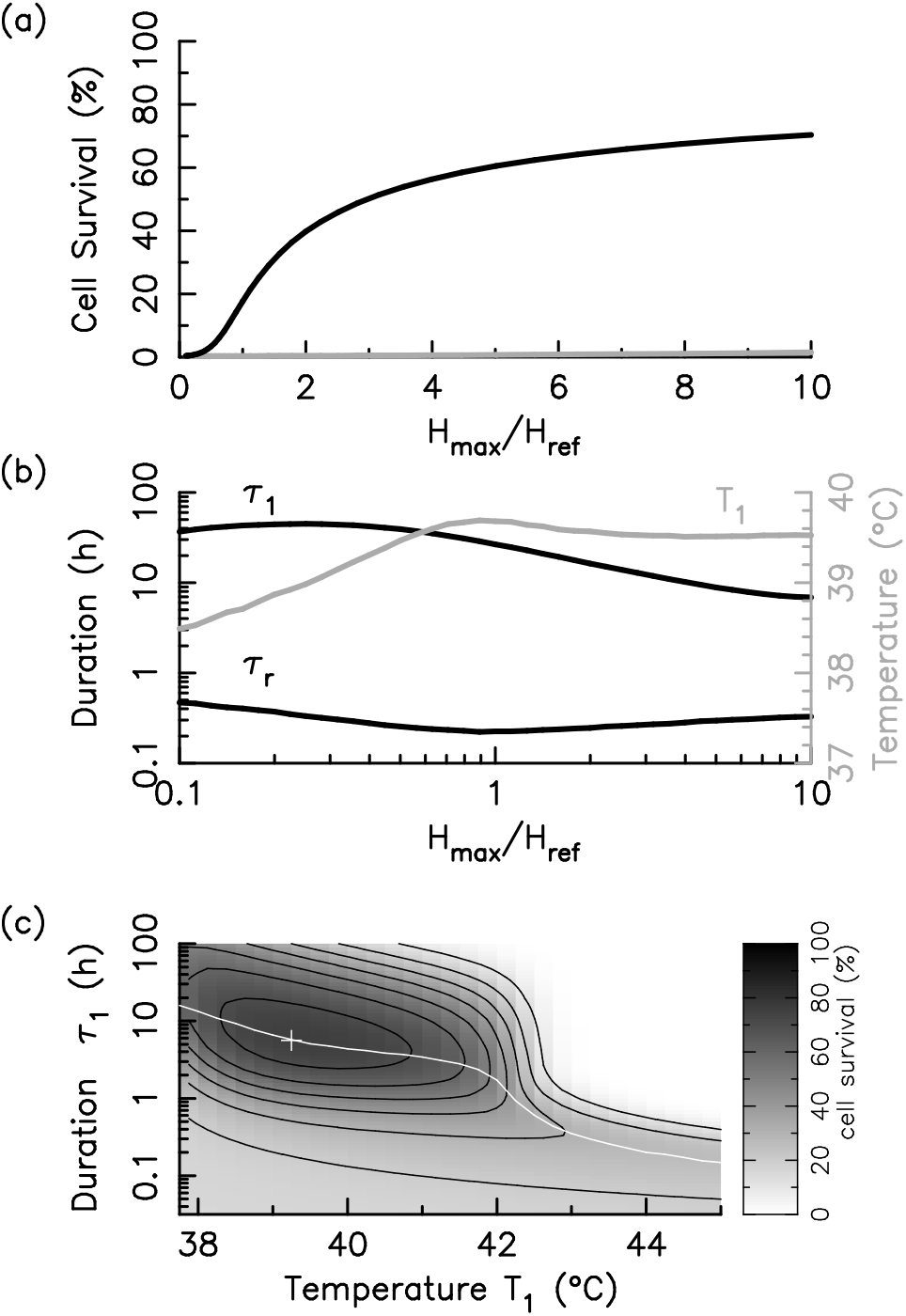
Optimized protocols inducing thermotolerance. The lethal hyperthermia protocol of reference is a temperature *T*_2_ =43°C during *τ*_2_ = 2 h with a chaperone capacity *H*_ref_ = 0.4 *μ*M (a) Cell survival without (grey line) and with (black line) priming. (b) Optimal parameters *τ_r_*, *T*_1_ and *τ*_1_ of the priming protocol. (c) Cell survival in the (*T*_1_, *τ*_1_) plan for *H*_max_ = 10 *H*_ref_ and *τ*_2_ = 2 h; the isovalues 10%, 20%, 30%, 40%, 50%, 60%, 70% (solid black lines); optimal manifold (solid white line) and the maximum value (white cross).

## 3. Discussion

In the present study, a simple model of biochemical adaptation and cell survival upon heat stress provides mechanistic and quantitative understanding of stress priming in the context of thermotolerance. The model is based on and extend previous data-driven modeling investigations of transcriptional control of chaperone synthesis [31, 32] and survival response to heat shock [27, 28]. Accordingly, the model displays a good qualitative agreement with a broad set of experiments while retaining a few number of free parameters which are likely to represent the diversity of cellular phenotypes including normal and cancer cells [28]. This framework is valuable to describe how priming or preconditioning stimulus dynamically modulates the intracellular state, but, more powerfully, to relate the characteristics of priming protocol and the survival benefit of induced thermoprotection and to optimize, in a cell type-specific manner, priming protocol design.

In the context of hyperthermia, an acknowledged feature of this improved tolerance is the flattening observed either by exposure to a long stress duration [36], or on particularly reactive cell lines such as CHO [37, 26] and colon cancer cells (PC3, RWPE) [38]. The standard thermal dosimetry technique, CEM43, does not predict nor explain such an effect, which is troublesome in the case of colon cancer since hyperthermia is one of the techniques used in the arsenal of possible treatments, notably to prevent recurrences in the intraperitoneal cavity in HIPEC-type protocols (Hyperthermic intra-peritoneal chemotherapy) [39]. The model reproduces several observed characteristics of thermotolerance such as the 1-hour temperature window, mainly between 41 and 43°C and the 4-hour recovery time to elicit significant thermotolerance effect [40, 22], which are explained by the saturation and timescale characteristics of the negative feedback loops by why damage activates chaperone synthesis with a maximal renaturation capacity.

Establishing thermotolerance characteristics for each cell line requires a vast number of experiments, and protocols were not always standardized, thus providing scattered data. This obviously complicates the implementation of new hyperthermia protocols relying insteady on trial-and-error approach. The here proposed modeling framework combines coarse-grained description, mechanistic understanding and predictive capacities, which only requires few measurements to be applicable to any cell types. A single cell survival assay for one exposure and a single measurement of chaperone relative abundance, before and after stress, are indeed sufficient to calibrate model parameters. In turn, model can predict the broad set of adaptation behaviors and fate output upon any thermal protocol.

The preconditioning/priming treatment can be used not only to modulate survival response to heat shock but also to control the synthesis and abundance of molecular chaperones in broader contexts. Indeed, molecular chaperones have other cytoprotective roles than protein folding, such as their role in anti-inflammatory processes [41]. Thus, a proposed strategy to fight against Acute Respiratory Distress Syndrome (ARDS) consists in increasing the level of molecular chaperone [42, 43], and it has recently been proposed to use hyperthermia to achieve this in chemical-free treatment for fragile patients [44]. This simple and inexpensive strategy were also proposed in the context of COVID19 crisis because clinical data indicate that severe COVID-19 most commonly manifests as viral pneumonia-induced acute respiratory distress syndrome (ARDS) [45]. A good knowledge of the adaptation mechanism would allow the optimization of the thermal protocol to maximize the abundance of molecular chaperones while maintaining control over cell survival (obviously targeting very high values in this case). In this sense also, the model we propose here could be skillfully used to propose new protocols.

## Acknowledgements

This work has been supported by the LABEX CEMPI (ANR-11-LABX-0007) and by the Ministry of Higher Education and Research, Hauts de France council and European Regional Development Fund (ERDF) through the Contrat de Projets Etat-Region (CPER Photonics for Society P4S).

